# PL-PatchSurfer3: Improved Structure-Based Virtual Screening for Structure Variation Using 3D Zernike Descriptors

**DOI:** 10.1101/2024.02.22.581511

**Authors:** Woong-Hee Shin, Daisuke Kihara

**Author notes:** **Correspondence to** Daisuke Kihara, Woong-Hee Shin.

## Abstract

Structure-based virtual screening (SBVS) is a widely used method in silico drug discovery. It necessitates a receptor structure or binding site to predict the binding pose and fitness of a ligand. Therefore, the performance of the SBVS is affected by the protein conformation. The most frequently used method in SBVS is the protein-ligand docking program, which utilizes atomic distance-based scoring functions. Hence, they are highly prone to sensitivity towards variation in receptor structure, and it is reported that the conformational change significantly drops the performance of the docking program.

To address the problem, we have introduced a novel program of SBVS, named PL-PatchSurfer. This program makes use of molecular surface patches and the Zernike descriptor. The surfaces of the pocket and ligand are segmented into several patches by the program. These patches are then mapped with physico-chemical properties such as shape and electrostatic potential before being converted into the Zernike descriptor, which is rotationally invariant. A complementarity between the protein and the ligand is assessed by comparing the descriptors and geometric distribution of the patches in the molecules. A benchmarking study showed that PL-PatchSurfer2 was able to screen active molecules regardless of the receptor structure change with fast speed. However, the program could not achieve high performance for the targets that the hydrogen bonding feature is important such as nuclear hormone receptors.

In this paper, we present the newer version of PL-PatchSurfer, PL-PatchSurfer3, which incorporates two new features: a change in the definition of hydrogen bond complementarity and consideration of visibility that contains curvature information of a patch. Our evaluation demonstrates that the new program outperforms its predecessor and other SBVS methods while retaining its characteristic tolerance to receptor structure changes. Interested individuals can access the program at kiharalab.org/plps3.

## INTRODUCTION

Virtual screening (VS) is a computational approach utilized in Silico drug discovery to identify potential active chemicals for a specific target protein. This method explores a library of molecules to search for promising candidates [1–3]. Within virtual screening, two main categories can be distinguished: ligand-based virtual screening (LBVS) and structure-based virtual screening (SBVS). In LBVS, the ligands in a library are evaluated by comparing them to a template active molecule [4–5]. LBVS methods are generally efficient and require only template compounds. However, their reliance on templates limits their ability to discover novel scaffold compounds since they do not consider receptor structures.

Conversely, SBVS employs different strategies such as protein-ligand docking and receptor pharmacophore search to predict the binding position of a ligand within the target receptor pocket and determine the binding affinity [6–8]. Unlike LBVS, SBVS utilizes the receptor structure and does not rely on prior knowledge of known drugs. As a result, the hits obtained from a ligand library through SBVS tend to be more diverse compared to LBVS.

The majority of SBVS methods, particularly docking programs, rely on an atomic description to calculate the binding energy. In this approach, the Gibbs binding free energy (ΔG_bind_) of a protein-ligand complex structure is determined by the sum of atomic pair interaction energy, which is dependent on the distance between atoms. Although this method has gained widespread use, it possesses certain limitations. One primary concern is its susceptibility to changes in the receptor conformation. The atom pair interaction function penalizes close proximity between atoms, and this is particularly evident in force-field-based scoring functions where the repulsive term of van der Waals energy follows a 1/r^12^ function. Consequently, even minor alterations in the receptor structure, such as a tautomeric state, can greatly impact the results and lead to failures, particularly in homology-modeled structures [9,10]. Another limitation of SBVS methods is the computational speed. In order to identify the optimal docking pose that minimizes ΔG_bind_, the docking program needs to perform numerous energy calculations. To expedite the docking process, many SBVS methods employ a grid consisting of pre-computed energies and utilize trilinear interpolation. Despite these optimizations, the conventional protein-ligand docking program still requires several minutes to dock a single molecule using a single CPU. Consequently, this approach may not be suitable for ultra-large-scale virtual screening scenarios involving the search of millions of compounds [11,12]. The extensive computational time involved poses a significant challenge in conducting efficient and timely screenings at such scales. Thus, an ideal virtual screening method would combine the advantages of both approaches: the ability to quickly discover diverse active compounds and the applicability to target receptors without experimentally solved structures.

In 2016, we introduced a novel SBVS program called PL-PatchSurfer2 [10] as a solution to address the limitations of traditional methods. This program utilizes a surface patch description to capture the complementarity between a protein and a ligand, instead of relying on atomic details. The binding pocket of the receptor and the ligand are segmented into a number of overlapping patches, and four physicochemical characteristics (shape, electrostatic potential, hydrogen bonding, and hydrophobicity) are assigned to each patch. These characteristics are then transformed into a three-dimensional Zernike descriptor (3DZD) [13,14], which is the coefficients of a series expansion using Zernike basis functions. The rotational invariance feature of the descriptor enables the comparison of 3D shapes without the need for alignment. This approach has been successfully applied to various biological problems, including protein-protein docking [15–17], protein-protein interface comparison [18], ligand-ligand comparison [19], pocket-pocket comparison [20,21], electron density maps from electron microscopy [22,23], and protein structure similarity search [24,25].

By incorporating the 3DZD, PL-PatchSurfer2 offers advantages in terms of tolerance to receptor conformational changes and computational speed. Benchmark results have demonstrated that our program maintains its performance on homology-modeled structures and apo structures, achieving an enrichment factor (EF) at ∼ 80% of the holo structure performance. In contrast, other SBVS programs exhibited significant deterioration when applied to these structures [10]. This highlights the effectiveness and robustness of PL-PatchSurfer2 in virtual screening applications.

In this paper, we introduce a newer version of our SBVS program called PL-PatchSurfer3 (PL-PS3), which builds upon the foundations of PL-PatchSurfer2 while incorporating important updates. PL-PS3 brings about one change and one additional feature compared to its predecessor. The change involves the hydrogen bonding term, which is modified from being represented by the 3DZD descriptor to utilizing surface point differences. In PL-PatchSurfer2, the 3DZD hydrogen bonding term is sometimes zeroed out due to most surface points having a neutral hydrogen bonding character. Consequently, the program fails to capture the interaction through hydrogen bonding between a protein and a ligand properly, resulting in zero enrichment factor (EF) at 1% for targets such as estrogen receptor alpha [10]. To address this limitation, PL-PS3 changes the definition of hydrogen bonding complementarity that takes into account the hydrogen bonding through surface point differences. The new program also introduces a new feature called visibility to better describe the characteristics of a molecular surface. The visibility feature takes into account the curvature of a surface patch, which cannot be adequately captured by the shape feature alone. By incorporating the visibility feature to account for surface curvature and updating the hydrogen bonding term, PL-PS3 offers a more comprehensive representation of molecular surface characteristics. These updates contribute to the program’s ability to accurately capture important molecular interactions and improve its performance in virtual screening tasks. The performance of PL-PS3 is benchmarked to Directory of Useful Decoys (DUD) set [26] first and compared to the conventional SBVS methods, AutoDock Vina [27] and DOCK6 [28]. Our program performed better than or comparable to the other SBVS methods. The programs were further tested on the apo structures. Finally, we screened the molecules to the model structures generated by MODELLER [29] and AlphaFold2 [30]. Our result showed that PL-PS3 outperforms other programs, especially for the apo and modeled structures.

## METHODS

### The Algorithm of PL-PatchSurfer3

The original algorithm of PL-PS3 was described in our previous publications [10,31]. In this section, we briefly introduce the algorithm focusing on the newly implemented/modified features in the program. The overall workflow of the program is illustrated in Figure 1.

**Figure 1.**
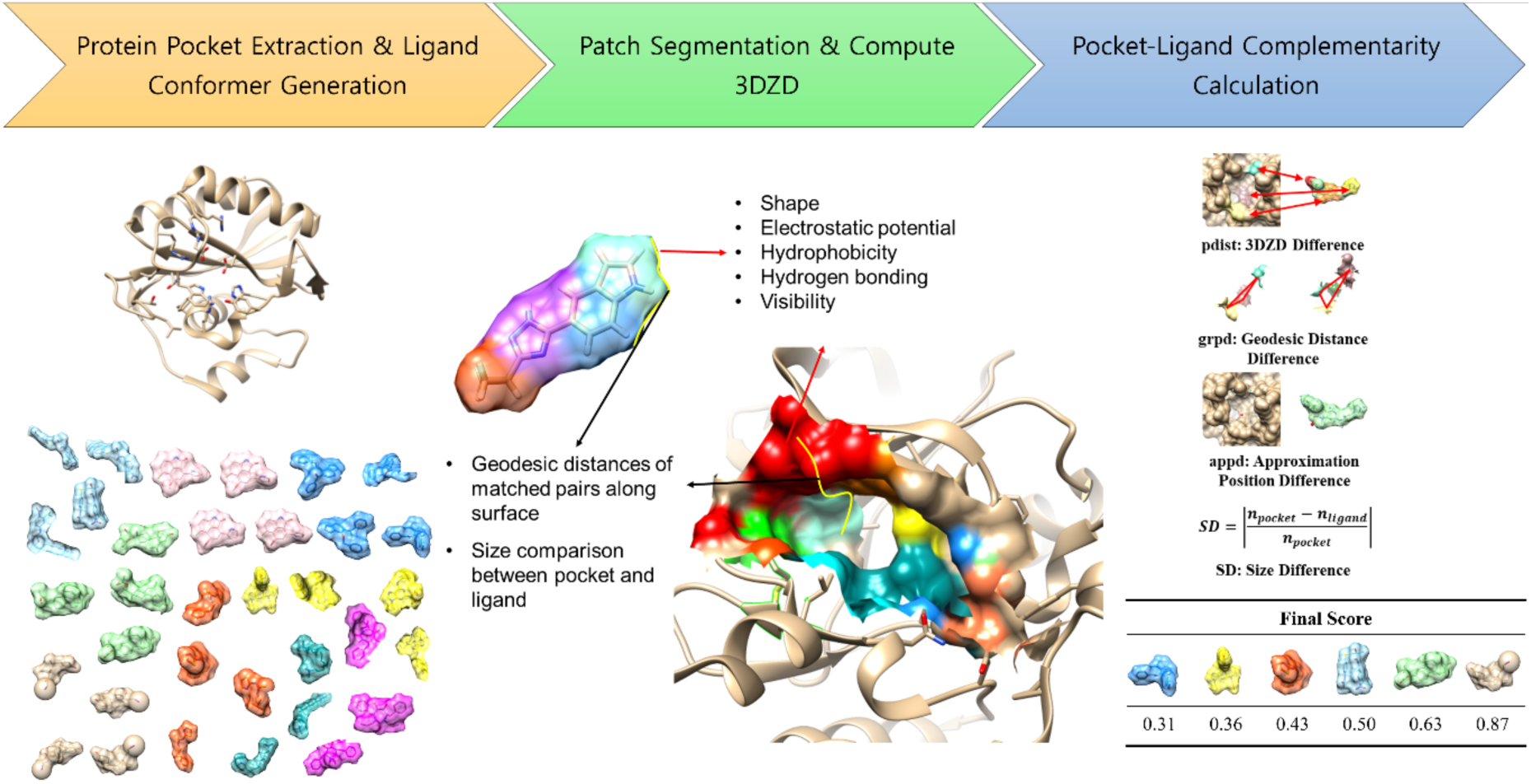
The overall workflow of PL-PatchSurfer3.

### Generating Patches and Calculating 3DZD

The objective of PL-PS3 is to evaluate the complementarity between a given binding pocket and a ligand. To account for the flexibility of the ligand, a pre-generated set of 3D conformations is obtained by conformer generation programs such as OMEGA [32]. The molecular surfaces and electrostatic potential on the surfaces of both the binding pocket and ligand conformations are generated using Adaptive Poisson-Boltzmann Solver (APBS) [33]. Once the molecular surface is obtained, PL-PS3 proceeds to calculate the hydrophobicity, hydrogen bonding, and visibility of the surface.

The novel addition to PL-PS3 is the visibility feature, which focuses on the local curvature of surface points [34] and was not fully captured by the shape feature in the previous version. The visibility calculation in our program involves casting rays from the surface point of interest to its neighboring voxel points. In PL-PS3, a total of 512 rays are cast from each surface point. If a neighboring point is a surface point, the corresponding ray is considered blocked. The visibility value is then determined by dividing the number of unblocked rays by the total of 512 rays. As a result, the visibility score ranges from 0, indicating a fully buried point, to 1, representing a fully exposed point.

Following the assignment of all characteristics to the surface points, the surface of the given molecule conformation is divided into overlapped patches, starting from selecting seed points, a center of patches. The minimum distance of 3 Å is set between seed points and spheres with a radius of 5 Å are positioned at each seed point to define the boundaries of the individual patches.

Each patch in PL-PS3 is characterized by five physicochemical features: shape, electrostatic potential, hydrophobicity, visibility, and hydrogen bonding. Except for hydrogen bonding, these features on the surface patch are converted to a 3DZD using Equation (Eq. 1) [13,14].

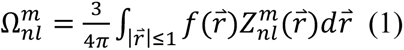

In Eq. 1, 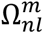 represents the 3D Zernike moments, 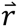 denotes the coordinates in three-dimensional space, 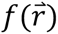 represents a scalar function in 3D space, and 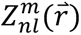 represents the Zernike-Canterakis basis function. The Zernike-Canterakis basis function is a product of the radial function *R_nl_*(*r*) and spherical harmonics 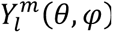, as described in Equation (Eq. 2).

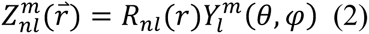

The Zernike-Canterakis basis function is determined by three integers: n, 1, and m. Similar to the wave function of a rigid rotor, these integers have specific conditions: -1 < m < l, 0 ≤ 1 ≤ n, and (n - 1) is an even number.

The 3DZD is finally obtained by normalizing 3D Zernike moments (Eq. 3).

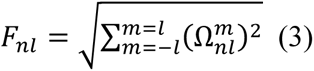

The dimension of 3DZD is determined by the order n. By setting n = 15, PL-PS3 has 72-dimensional vector.

### Paring Patches of a Protein Pocket and a Ligand

To determine the optimal patch pairs between a protein (P) and a ligand conformer (L), PL-PS3 utilizes a modified Auction algorithm, which is a computational optimization algorithm [21]. This algorithm aims to minimize the Patch Score by iteratively searching for suitable patch pairs. The Patch Score, calculated according to Equation (Eq. 4), serves as an objective function to guide the algorithm in its search for optimal patch pairs.

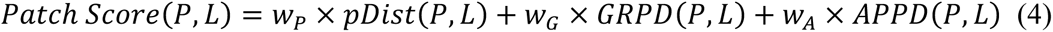

P and L represent pocket and ligand, respectively.

pDist, the first term, is a weighted sum of the distance between the matched patches. (Eq. 5)

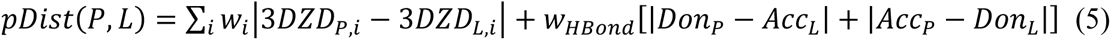

The index i represents the physicochemical features, including shape, electrostatic potential, visibility, and hydrophobicity. The distances for these features are calculated using the Euclidean distance between their 3DZDs. The hydrogen bonding component of pDist accounts for the local nature of this feature and is treated differently from the other components. Instead of using the 3DZD distance, the hydrogen bonding distance is calculated by summing the differences between the number of surface points in the protein pocket with donor character (Don_P_) and acceptor character in the ligand (Acc_L_), as well as the number of surface points in the ligand with donor character (Don_L_) and acceptor character in the protein (Acc_P_). The weights were trained on the PDBBind core set [35] to find the interacting patches between a protein and a ligand. The interacting patch pair is defined as the distance between their seed points shorter than 3.0 Å in the crystal structure. The optimized weight values are 1.0, 0.8, 1.4, 0.8, and 0.1 for shape, electrostatic potential, visibility, hydrogen bonding, and hydrophobicity, respectively.

Geodesic Relative Patch Difference (GRPD) and Approximate Patch Position Difference (APPD) terms play a role in evaluating and optimizing the placement of matched patch pairs based on geodesic distances. These terms are essential for accurately locating the patch pairs within the pocket and ligand. The GRPD term quantifies the difference in geodesic distances between the matched patch pairs and other patch pairs. By minimizing the GRPD, the algorithm ensures that the matched patch pairs are appropriately positioned relative to other patch pairs in both the pocket and the ligand. On the other hand, the APPD term involves the construction of a histogram of geodesic distances, Approximate Patch Position (APP), from a seed point to other points on the surface. This histogram is created using a bin size of 1 Å, allowing the approximate position of a patch within the molecule to be represented by a vector. The APPD is then calculated using Equation (Eq. 6), which captures the difference in the approximate patch positions between the pocket and the ligand.

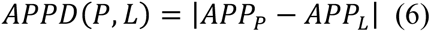

After finding the optimal patch pairs, the complementarity between a given pocket and a ligand conformer is obtained by the sum of three terms (Eq. 7). The weights for Patch Score (Eq. 4) are also trained on the PDBBind set, yielding 0.6, 0.4 and 1.05 for pDist, GRPD, and APPD, respectively.

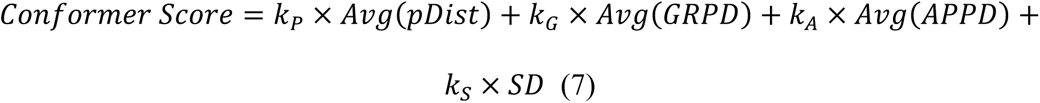

The Avg(X) is an average of a term X for all matched pairs. The last term, SD is a size difference between the pocket and the ligand conformer, calculated by the difference in the number of patches of given molecules. All weights (k_i_) were optimized to maximize the benchmark performance.

### Scoring Functions Used for Ranking Compounds

Since a compound may have up to 50 conformations, it is necessary to select a representative score to rank the molecules in the library and identify the active compounds. In PL-PatchSurfer, two types of scoring functions were utilized. The first function is the Lowest Conformer Score (LCS). This function determines the lowest Confomer Score (Eq. 7) among the conformations, hence providing the representative score of the compound. The Boltzmann-Weighted Score (BWS) is an alternative method of scoring that uses a weighted average of all conformations instead of relying on a single conformation score (Eq. 8) as below.

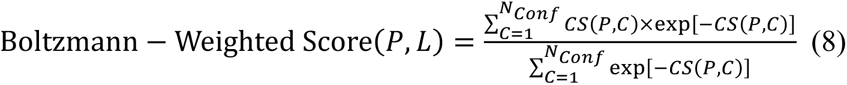

The score is calculated by taking the sum of each Conformer Score (CS) between the protein and each conformation generated by OMEGA, multiplied by the exponential of negative CS(P,C), divided by the sum of the exponential of negative CS(P,C) values across all conformations (N_Conf_). In Equation 8, P represents the protein, L represents the ligand, and C represents each generated conformation.

### Benchmark Datasets

To evaluate the performance of PL-PS3, the DUD dataset was primarily used [26]. The DUD dataset consists of 40 proteins, each with a ligand library composed of active compounds that have been experimentally validated and decoy compounds that possess similar chemical features. This dataset provides a reliable benchmark for evaluating virtual screening methods. For the evaluation, the same benchmark set as PL-PatchSurfer2 was employed. Specifically, 15 targets that could not be processed by APBS were excluded from the evaluation. The ratio between active compounds and decoy compounds in the dataset was set to 1:29, ensuring a realistic representation of the screening scenario. We used the holo structures that were provided in the DUD dataset.

In addition to evaluating PL-PS3 on holo structures, we also conducted tests on apo structures. In the apo form, the protein lacks a bound ligand and may exhibit different conformational states compared to the holo form. It is known that the performance of SBVS programs is generally lower on apo structures, as the absence of a ligand can affect the accuracy of the predictions. We gathered 18 apo structures available from Protein Data Bank [36] for evaluation. By including apo structures in the assessment, we aimed to assess the robustness and performance of PL-PS3 in situations where the protein is not bound to a ligand. This allows us to understand the program’s capability to handle diverse protein conformations and identify potential ligand binding sites even in the absence of a bound ligand.

The lack of experimentally determined protein structures poses a significant challenge for SBVS, as the availability of a receptor structure is crucial for accurate predictions. Homology modeling is a commonly used technique to predict protein structures based on their sequence similarity to known structures. However, SBVS programs have shown poor performance on homology modeled structures [9,10]. To address this challenge, we evaluated the performance of PL-PS3 and other SBVS methods on a benchmark set of modeled receptor structures. The benchmark set was generated using MODELLER [29] to perform homology modeling VS. In addition, recent advances in protein modeling using deep-learning techniques such as AlphaFold2 [30] and RoseTTAFold [37] have shown remarkable accuracy in predicting protein structures. Therefore, we downloaded AlphaFold2 models from KinCoRe [38]. Dunbrack and his colleagues constructed catalytic domains of human kinases using AlphaFold2 [39]. They curated multiple sequence alignment to model the active state of kinases. Six kinase structures (CDK2, EGFr, FGFr1, P38 MAP, SRC, and VEGFr2) were used to investigate whether the SBVS performances of the benchmarked programs depend on the quality of models or not.

### Evaluation Metrics

To assess the performance of the SBVS methods, several evaluation metrics were used, including enrichment factor (EF), area under the receiver operating characteristic curve (AUC), and Boltzmann-enhanced discrimination of receiver operating characteristic (BEDROC).

The EF measures the ability of a method to selectively retrieve active compounds compared to decoy compounds. EF at various percentages (1%, 5%, and 10%) represents the ratio of the percentage of retrieved actives to the expected percentage based on random selection.

The AUC is a widely used metric that evaluates the overall discriminatory power of a method. It measures the ability of the method to rank active compounds higher than decoy compounds across a range of classification thresholds. A higher AUC value indicates better performance.

To address the early recognition problem associated with AUC [40], BEDROC is used as an alternative metric. BEDROC incorporates a Boltzmann weight factor that assigns higher weights to early-ranked actives, giving more importance to early recognition. This metric provides a more focused evaluation of early enrichment performance.

### Protocols for Compared SBVS Programs

In the comparative evaluation, PL-PS3 was benchmarked against PL-PatchSurfer2 as well as popular SBVS methods, namely AutoDock Vina [27] and DOCK6 [28]. To conduct the SBVS experiments, the first step was to define the binding pocket for each method. The pockets of the holo structures were defined using the co-crystalized ligands present in the crystal structure. For apo and modeled structures, the binding pockets were defined same the holo structures, after aligning the structures using TM-align [41] and copy the ligands. The protonation states and atomic charges of the structures were assigned using UCSF Chimera [42].

For AutoDock Vina, the docking box was defined as a cube with dimensions of 25 Å on each side. For DOCK6, the docking box was constructed with a 5 Å margin from the space occupied by the cognate ligand. The other parameters of the three programs were set to their default values.

## RESULTS AND DISCUSSION

### Training and Benchmark on holo structure set

To determine whether the compound is complementarity to the binding pocket or not, the weights in the Eq. 7 should be determined. The 25 targets of the DUD set are divided into two sets, the same as in the previous paper [10]. The list of proteins for each set is listed in Supplementary Information Table S1. The weights were optimized to get high BEDROC values for each set and the determined weights were tested on the other protein set. The optimized weights for set 1 and set 2 are (k_p_=0.2, k_G_=0.4, k_A_=1.0, k_S_=0.7) and (k_p_=0.4, k_G_=0.8, k_A_=0.7, k_S_=1.0), respectively. The test result is summarized in Table 1. Individual protein results are given in Supplementary Information Table S2.

**Table 1.** Holo screening result. The results of PL-PS3 are the average of test results. The highest value in each metric among the benchmarked programs is highlighted as bold.

| Program | EF1% | EF5% | EF10% | AUC | BEDROC |
| --- | --- | --- | --- | --- | --- |
| PL-PS3 (BWS) | 14.35 | **5.21** | 3.34 | 0.606 | **0.313** |
| PL-PS3 (LCS) | 13.91 | 4.90 | 3.10 | 0.567 | 0.291 |
| PL-PatchSurfer2 (BWS) | **14.79** | 5.18 | 3.09 | 0.577 | 0.308 |
| PL-PatchSurfer2 (LCS) | 13.74 | 4.71 | 2.84 | 0.539 | 0.280 |
| AutoDock Vina | 7.92 | 5.05 | **3.37** | **0.632** | 0.276 |
| DOCK6 | 11.47 | 4.02 | 2.47 | 0.502 | 0.239 |

Compared with PL-PatchSurfer2, the new version showed improved performance except for EF1% of BWS scoring. The changes from PL-PatchSurfer2 are changing of hydrogen bonding term and adding visibility. To incorporate the local feature of hydrogen bonding, PL-PS3 calculates differences in surface points instead of the Euclidean distance of 3DZDs. This modification improves the performance of the targets with hydrogen bonding is important. For example, with LCS scheme, EF1% of ERα agonist and antagonist are improved from 9.0 to 13.50 and 0.0 to 8.18, respectively. The binding site of the nuclear hormone receptor family has common features: a hydrophobic region located in the middle, while the two ends have hydrogen bonding functional groups making a hydrogen bonding network with water and ligand molecules [43]. Comparing the scoring schemes of PL-PS3, BWS scoring showed higher performance in all metrics than LCS, which is consistent with the previous studies [10]. When comparing PL-PS3 with traditional SBVS programs such as DOCK6 and AutoDock Vina, the results revealed differences in their performance across different evaluation metrics. AutoDock Vina demonstrated the highest performance in terms of EF at 10% and AUC. On the other hand, PL-PS3 exhibited the best performance in EF at 1%, 5% and BEDROC.

Since the protein is a flexible molecule, the binding site adopts the conformation that fits to its partner. Thus, static picture of SBVS during screening could lead the bias in high-ranked compounds towards the molecules similar to the co-crystalized one. To investigate patch description might avoid the bias or not, we divided 25 targets into four, based on the average dissimilarity of active compounds from the co-crystalized molecule. SIMCOMP [44] was used to calculate the similarity between compounds based on 2D structure, and dissimilarity was computed as (1 – similarity). Table 2 shows the average EF1% values of benchmarked programs.

**Table 2.** Average EF1% of the divided set based on average 2D dissimilarity of active compounds from the co-crystalized molecule. The numbers in the parentheses are the number of proteins. The best value in each set is highlighted as bold.

| Average Dissimilarity | PL-PS3 (BWS) | PL-PS3 (LCS) | AutoDock Vina | DOCK6 |
| --- | --- | --- | --- | --- |
| > 0.80 (5) | **18.78** | 17.20 | 4.00 | 8.80 |
| > 0.70 (10) | **14.45** | 14.22 | 6.81 | 12.77 |
| > 0.60 (3) | **15.00** | 13.41 | 5.40 | 9.52 |
| < 0.60 (7) | 10.78 | 11.35 | **13.37** | 12.34 |

For the 18 targets where dissimilarity exceeded 0.6, PL-PS3 outperformed traditional SBVS methods. However, for targets with dissimilarity levels below 0.6, our program demonstrated lower EF1% compared to traditional methods. Notably, when the dissimilarity surpassed the cutoff point of 0.6, AutoDock Vina showed a 50% decrease in EF1%, and DOCK6 performed poorest for proteins with dissimilarity above 0.8. On the other hand, PL-PS3 tends to perform consistently. This could potentially be an advantage of utilizing patch-based description, as it discovers diverse compounds in high-rank by finding key patches regardless of the geometrical similarity of the compounds. Re-optimizing the Conformer Score weights using the seven targets with dissimilarity < 0.6 puts high weights on the geometrical components (k_p_=0.1, k_G_=1.0, k_A_=0.6, k_S_=1.0), making EF1% as 13.58. This means that global shape is important for the targets with low dissimilarity.

One of the targets that has distant active molecules from the co-crystalized compound is HIVPR. The average dissimilarity is 0.804. The EF1% values of PL-PS3 with both scoring schemes are 24 (BWS) and 20 (LCS), while those of AutoDock Vina and DOCK6 are 4 and 2, respectively. Figure 2 illustrates examples of compound that are not similar to the cognate ligand. The 1^st^ and 2^nd^ ranked molecules by PL-PS3 are ZINC3833854 and ZINC3833846 with dissimilarity from the co-crystalized compound 0.805 and 0.606, respectively. The compounds were ranked at 23^rd^ (ZINC3833854) and 22^nd^ (ZINC3833846) by AutoDock Vina out of 1590 compounds. DOCK6 placed the molecules 1548^th^ and 1439^th^, respectively.

**Figure 2.**
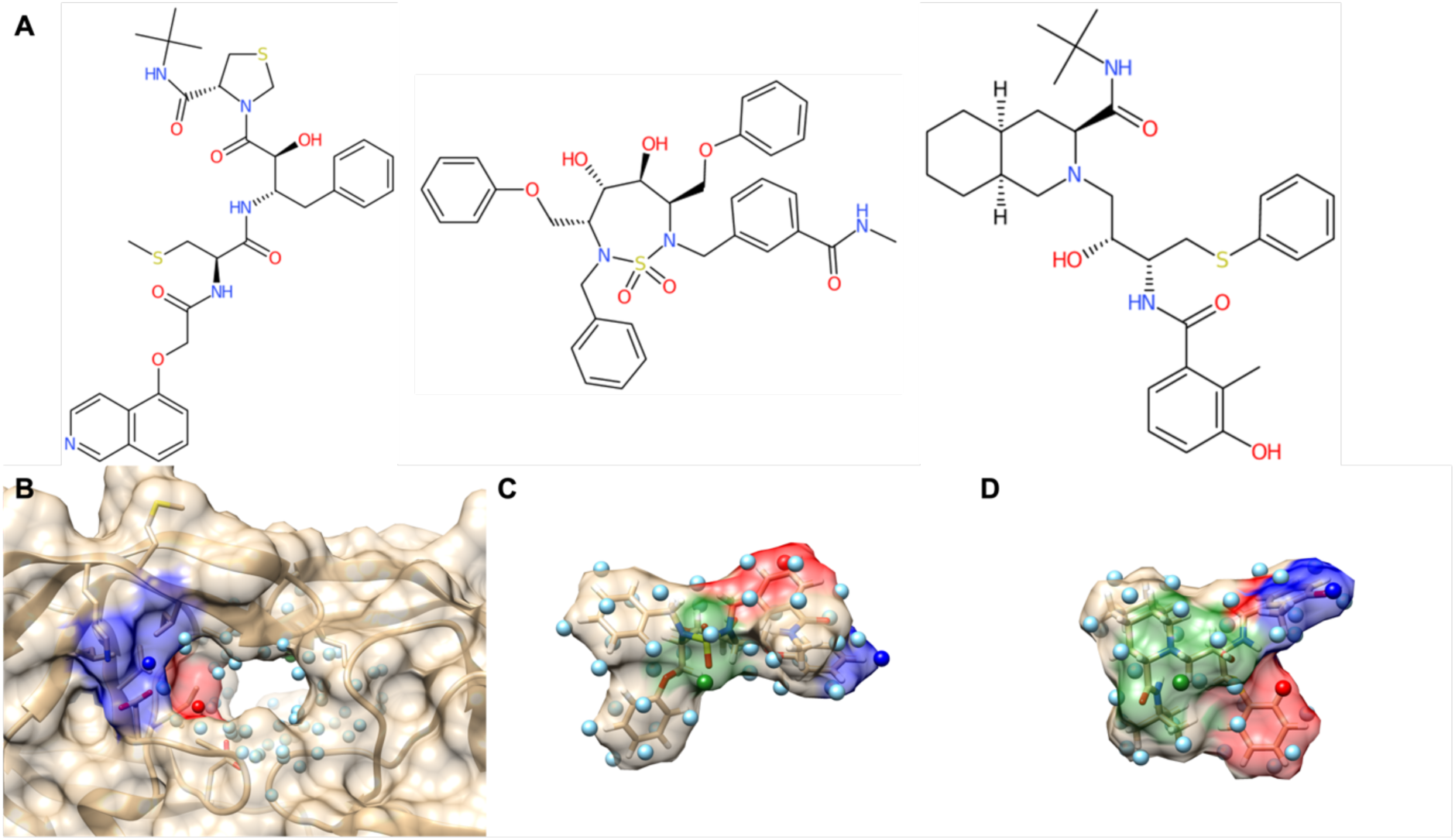
**A.** 2D structure of HIVPR co-crystalized compound (left), ZINC3833854 (middle), and ZINC3833846 (right). **B.** HIVPR binding pocket surface and seed points. **C. D.** ZINC3833854 and ZINC3833846 molecular surfaces and seed points. The seed points represented as spheres. The seed points of paired patches within the top 10 lowest score matched by PL-PatchSurfer3 have same color codes.

### Benchmark Result on Apo Structure Set

The flexibility of proteins introduces a challenge in SBVS, particularly when comparing the performance of methods on apo structures (ligand-absent) and holo structures (ligand-bound). The structural differences between these two forms might significantly impact the performance of traditional SBVS methods, often leading to a decrease in their performance.

Our previous study has shown that the structural change from holo to apo structures can result in a deterioration of over 30% in the performance of traditional SBVS methods, particularly in terms of EF at 1%. However, it was observed that PL-PatchSurfer2 managed to retain its performance in this scenario [10]. We tested our new program on the apo structures and obtained a similar result as the previous work. Table 3 summarizes the result for the apo set (Individual result is shown in Supplementary Information Table S3). The EF at 1% decreased from 13.71 to 13.16 (97%) for BWS scheme and from 13.02 to 10.75 (83% of the holo-structure) for LCS scheme. In addition, different from the holo-structure result, PL-PS3 showed the highest performance in all metrics.

**Table 3.**
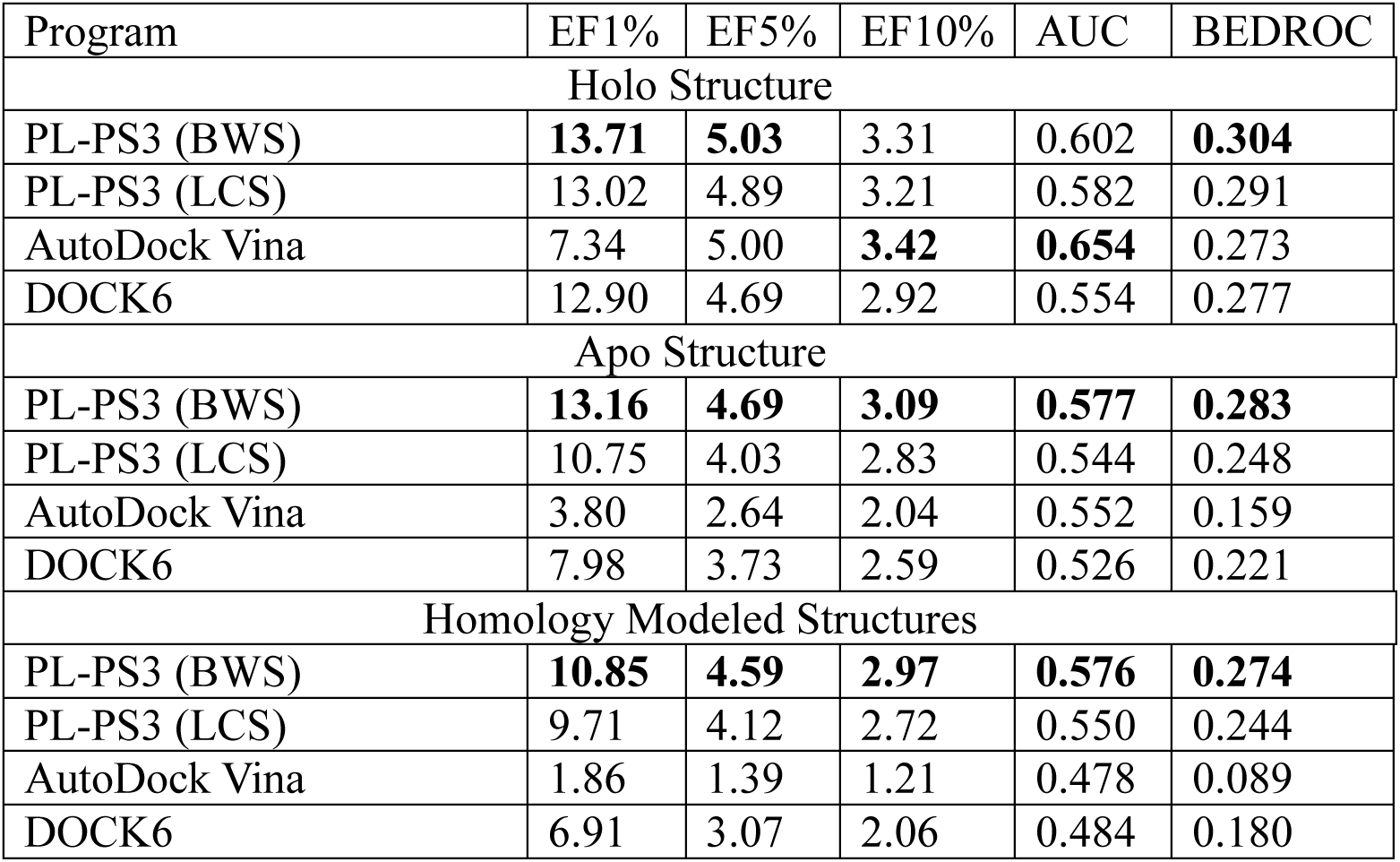
Performance of benchmarked proteins with receptor structure variation. The best values are highlighted as bold.

The ability of PL-PS3 to maintain its performance on apo structures suggests that its patch-based approach captures key interactions and complementarity between the protein and ligand, even in the absence of the bound ligand. By focusing on local patches and their physicochemical characteristics, PL-PS3 can potentially overcome the challenges associated with structural differences between apo and holo structures, enabling more reliable virtual screening results.

The impact of structural changes on the performance of traditional SBVS methods is evident in the case of the three proteins (RXRα, HIVPR, and SRC) with binding site Ca-RMSD larger than 2 Å among the 18 apo structures. For these three proteins, the EF at 1% values of AutoDock Vina and DOCK6 showed a significant decrease, dropping from 10.78 to 1.33 and from 18.44 to 1.67, respectively. This decrease in performance is much higher compared to the average values, indicating that the structural differences strongly affect the performance of traditional methods in these particular cases.

In contrast, PL-PS3 demonstrated more robust performance, with EF at 1% values of 19.67 (18.33) and 16.44 (14.56) of BWS (LCS) scheme for the holo and apo structures, respectively. This suggests that the local patch comparison approach employed by PL-PS3 is particularly suitable for proteins with highly flexible receptors, allowing it to better handle the structural changes between apo and holo-structures. These results highlight the advantages of the patch-based approach in PL-PS3, as it captures local interactions and complementarity, which can help overcome the challenges posed by structural variations.

### Benchmark Results on the Homology Modeled Structures

When the target of interest does not have an experimental structure, structure prediction is widely used for SBVS. One of the most widely used structure prediction programs was MODELLER [29]. Similar to the apo structure, it is reported that the modeled structure also deteriorates the performance of SBVS [9,10]. Thus, we tested PL-PS3 with the traditional methods to investigate whether our program can also retain the performance or not. The 18 protein structures were predicted using MODELLER. Table 3 shows the performance of three SBVS programs on the modeled structure and the holo-structures. The two traditional SBVS methods failed to rank the active compounds in a higher rank. Comparing EF at 1%, AutoDock Vina and DOCK6 dropped from 7.34 to 1.86 and 12.90 to 6.91, respectively. In contrast, PL-PS3 tends to maintain its performance, changing from 13.71 to 10.85 with BWS scheme. In addition, PL-PS3 has the highest average values in all metrics. Individual protein result is given in Supplementary Information Table S4. The quality of homology models depends on the sequence similarity between the query protein and the target structure. It is known that the proteins that share ∼30% of the sequence identity have a similar fold [45]. The range 25% - 30% is called ‘twilight zone’. Out of 18 modeled structures, half of them were built from the template with sequence identity above 30%. Table 4 summarizes the performances of SBVS programs depending on the sequence identity.

**Table 4.** Performance of SBVS programs on homology modeled set. The set is divided into two depending on the sequence identity between the query and template sequences. The highest values are highlighted as bold.

| Sequence Identity (%) | PL-PS3 (BWS) | PL-PS3 (LCS) | AutoDock Vina | DOCK6 |
| --- | --- | --- | --- | --- |
| EF1% |  |  |  |  |
| > 30% | 10.74 | 10.40 | 2.04 | **11.24** |
| = 30% | **10.96** | 9.03 | 1.69 | 2.58 |
| BEDROC |  |  |  |  |
| > 30% | 0.214 | 0.207 | 0.087 | **0.245** |
| = 30% | **0.334** | 0.282 | 0.092 | 0.116 |

Regardless of the sequence identities, AutoDock Vina had the worst performance among the programs. DOCK6 performed the best in both metrics for the nine receptor structures modeled from the template with a sequence identity greater than 30%. However, its performance significantly decreased when the receptor structures were modeled from a sequence identity of 30%. On the other hand, PL-PS3 demonstrated consistent results in EF1% and even improved in terms of BEDROC. Our program showed superior performance for models with lower sequence identities, which may have deteriorated structures, compared to traditional methods. Figure 3 showed an example of SRC kinase. The sequence identity between query and template sequences was 39%. Due to the absence of the co-crystalized ligand, the beta-strands of homology model move inward to the binding pocket, narrowing the binding pocket (Figure 3A). The enrichment factor of PL-PS3 on holo structure is 20.00. When the homology-modeled structure is used, EF1% is decreased to 15.00 with LCS scheme. On the other hand, DOCK6 is changed from 23.33 to 16.67 and AutoDock Vina is dropped from 3.33 to 0.00. Out of 1800 molecules screened, all programs ranked ZINC1916192 within top 10% using holo structure as the receptor structure. PL-PS3, AutoDock Vina, and DOCK6 ranked the molecule at 120^th^, 56^th^, and 7^th^, respectively. However, when the homology-modeled structure is used the compound is ranked 119^th^, 1522^nd^, and 1443^rd^ by PL-PS3, AutoDock Vina, and DOCK6, respectively. PL-PS3 tried to match the patch with similar residues although the receptor structure was changed (Figure 3B). The hydrophobic patch (red), which covers the center benzyl moiety, is matched to the LEU273 and MET283 of both receptor structures. Similarly, the green patch, composed of a benzene ring and sulfonyl group, tends to have interaction with SER345 and LEU393. The pink patch of an aromatic ring with two hydroxyl groups also matched to the same residues of the receptor structures, TYR340 and SER342. On the other hand, the conventional docking programs placed the ligand out of the binding pocket.

**Figure 3.**
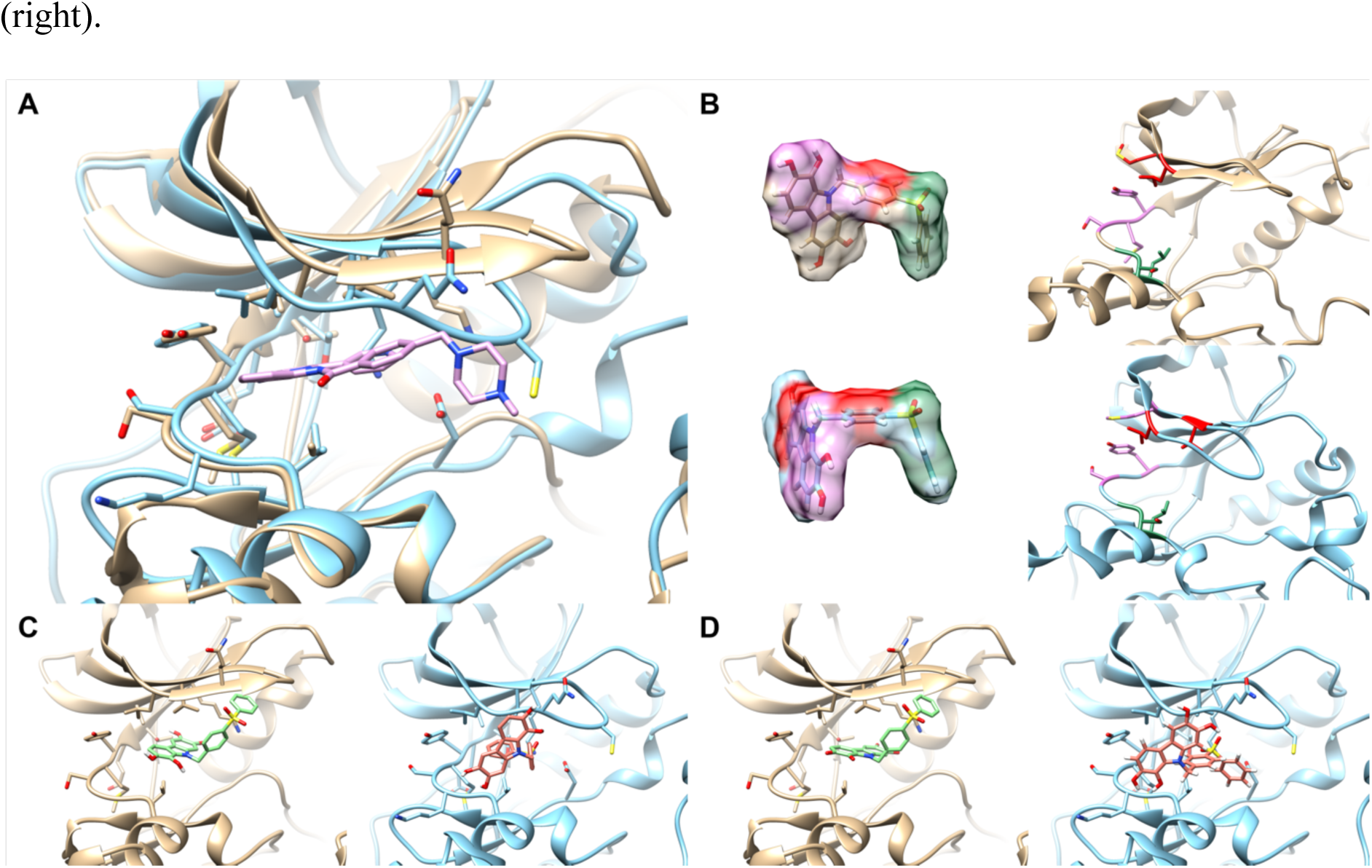
**A.** Binding pocket conformations of SRC holo structure (gold) and homology modeled structure (sky blue). The co-crystalized ligand of holo structure is shown in pink. **B.** Top scored conformation of ZINC1916192 screened against the holo-structure (upper left) and the homology modeled structure (lower left). The right panels show the protein structure and selected patches matched to the ligand patches. The matching pairs of the colored in same color code. **C.** Predicted binding poses of the compound by AutoDock Vina docked to the holo structure (left) and the homology modeled structure (right). **D.** Predicted binding poses of the compound by DOCK6 docked to the holo structure (left) and the homology modeled structure (right).

### Benchmark Results on the AlphaFold2 Structures

Recently, deep-learning-based prediction methods have revolutionized the structure prediction field, with an extremely high accuracy. Out of DUD benchmark set, kinases are selected because the active loop and the conformation of DFG motif vary depending on the binding ligand type [38]. We extracted six AlphaFold2 kinase structures from KinCoRe [38,39] database to study whether the SBVS techniques are suitable for the recent modeling programs or not. The average dissimilarity of active compound from the co-crystalized ligand of the six proteins is 0.746, slightly higher than average over targets (0.712).

On the AlphaFold2 targets, PL-PS3 showed the best performance among the benchmarked programs (Table 5, individual result is given in Supplementary Information Table S5) in all metrics. Compared to the holo structure, our program maintained a performance higher than 84% of the holo structure result in both scoring schemes. On the contrary, EF1% of DOCK6 and AutoDock Vina decreased to 66% when AF2 structures were used, which are higher than the case of homology modeled case.

**Table 5.**
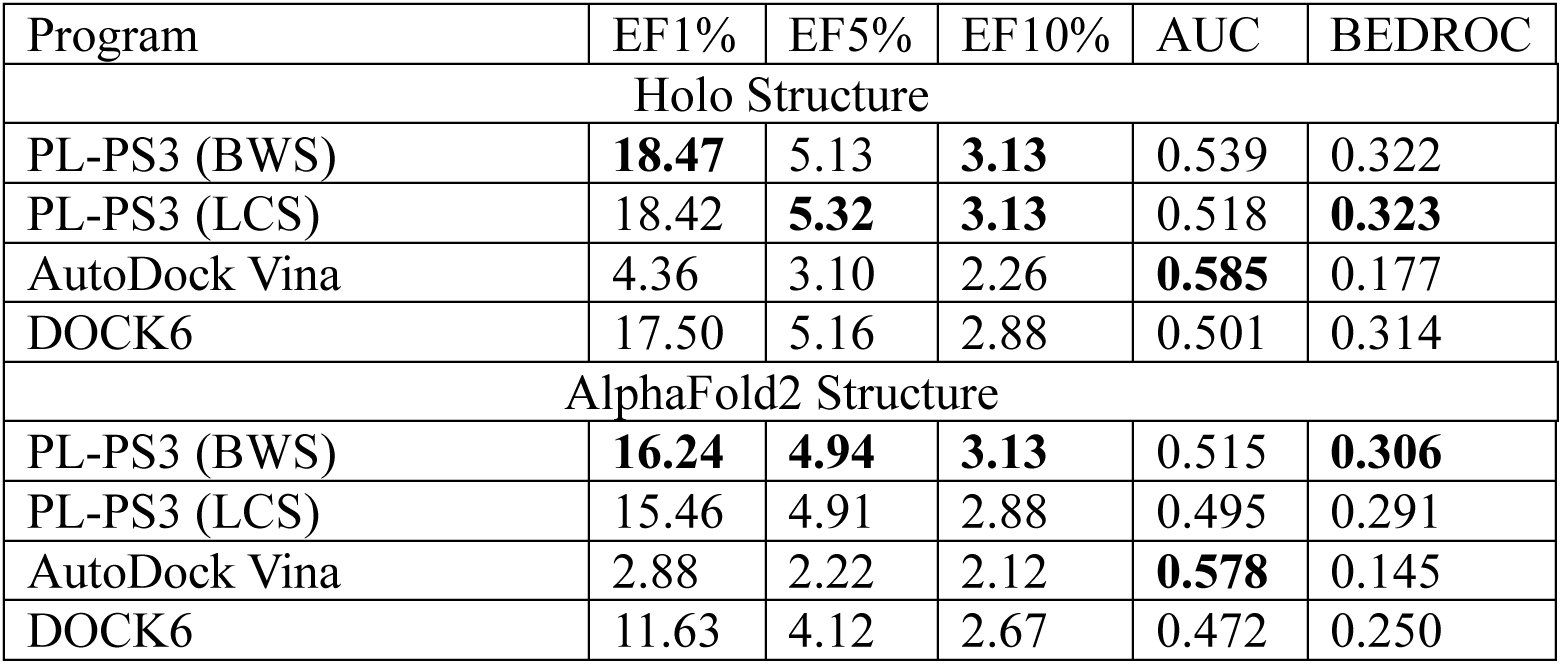
Benchmarking results of six kinases structures modeled by AlphaFold2. The highest values are highlighted as bold.

| Program | EF1% | EF5% | EF10% | AUC | BEDROC |
| --- | --- | --- | --- | --- | --- |
| Holo Structure |  |  |  |  |  |
| PL-PS3 (BWS) | **18.47** | 5.13 | **3.13** | 0.539 | 0.322 |
| PL-PS3 (LCS) | 18.42 | **5.32** | **3.13** | 0.518 | **0.323** |
| AutoDock Vina | 4.36 | 3.10 | 2.26 | **0.585** | 0.177 |
| DOCK6 | 17.50 | 5.16 | 2.88 | 0.501 | 0.314 |
| AlphaFold2 Structure |  |  |  |  |  |
| PL-PS3 (BWS) | **16.24** | **4.94** | **3.13** | 0.515 | **0.306** |
| PL-PS3 (LCS) | 15.46 | 4.91 | 2.88 | 0.495 | 0.291 |
| AutoDock Vina | 2.88 | 2.22 | 2.12 | **0.578** | 0.145 |
| DOCK6 | 11.63 | 4.12 | 2.67 | 0.472 | 0.250 |

## CONCLUSIONS

Structure flexibility is one of the key issues in SBVS. To overcome the obstacle, PL-PatchSurfer employed a local patch description instead of the atomic model used in conventional docking programs. Although our previous version showed comparable performance with the conventional programs, it has a problem of treating local features such as hydrogen bonding.

In the newer version, PL-PS3, instead of using Zernike descriptor, counting surface points with hydrogen bonding, and added visibility as a new physico-chemical property. With the help of the newly implemented terms, PL-PS3 showed better performance than its ancestor with improved results on nuclear receptors. In addition, the program maintains the advantages of its predecessor. PL-PS3 showed higher performance on the diverse compound set and non-holo receptor structures. These findings show that the distinct surface patch representation gives PL-PS3 an intriguing strength, namely enhanced tolerance to conformation change of targets. By offering more precise predictions in challenging situations, such as scaffold hopping or situations where only the apo form of targets or predicted structures of targets are accessible, PL-PS3 can significantly increase the capability of virtual screening by utilizing this special strength.

## Author Contributions

DK devised the study, while WHS conducted the benchmark, analyzed the outcomes, and composed the manuscript. DK revised and polished the article before its submission. All co-authors have duly reviewed the document.

## Availability of Data and Materials

The binary files and scripts of the program are available at kiharalab.org/plps3.

## List of Abbreviations

SBVS: Structure-Based Virtual Screening
VS: Virtual Screening
LBVS: Ligand-Based Virtual Screening
3DZD: Three-Dimensional Zernike descriptor
PL-PS3: PL-PatchSurfer3
EF: Enrichment Factor
DUD: Directory of Useful Decoys
APBS: Adaptive Possion-Boltzmann Solver
GRPD: Geodesic Relative Patch Difference
APPD: Approximate Patch Position Difference
APP: Approximate Patch Position
LCS: Lowest Conformer Score
BWS: Boltzmann-Weighted Score
AUC: Area Under the receiver operating characteristic Curve
BEDROC: Boltzmann-Enhanced Discrimination of Receiver Operating Characteristic

