## Supplementary Information Table S1 for "PL-PatchSurfer3: Improved Structure-Based Virtual Screening for Structure Variation Using 3D Zernike Descriptors"

**Table S1**. The target structures used for benchmark. The numbers in the parenthesis are the sequence identity between the holo structure and the templates.

| Set | Protein | Holo | Apo | TBM template |
|---|---|---|---|---|
| 1 | AmpC | 1XGJ | 1L0D | 3WRG (40%) |
| | AR | 1XQ2 | - | - |
| | CDK2 | 1CKP | 1HCL | 3VW6 (30%) |
| | COX-2 | 1CX2 | 5COX | 1EBV (64%) |
| | EGFr | 1M17 | 1M14 | 3BBT (78%) |
| | ER: antagonist | 3ERT | 2B23 | 4NQA (30%) |
| | FXa | 1F0R | 1C5M | 1SC8 (30%) |
| | HIVRT | 1RT1 | - | - |
| | PARP | 1EFY | 2PAW | 1GS0 (45%) |
| | PPARγ | 1FM9 | 1PRG | 1X7E (30%) |
| | RXRα | 1MVC | 1GLU | 1SJ0 (30%) |
| | VEGFr2 | 1VR2 | - | - |
| 2 | AChE | 1EVE | 1Q1H | 2OGS (30%) |
| | COX-1 | 1P4G | 1PRH | 1CVU (64%) |
| | ER: agonist | 1L2I | 2B23 | 1Z5X (30%) |
| | FGFr1 | 1AGW | 1FGK | 3PPZ (32%) |
| | GR | 1M2Z | - | - |
| | HIVPR | 1HPX | 3PHV | 3NR6 (30%) |
| | MR | 2AA2 | - | - |
| | NA | 1A4G | 1NSB | 1L7G (31%) |
| | P38 MAP | 1KV2 | 1P38 | 3OZ6 (35%) |
| | PDGFrb | model | - | - |
| | PR | 1SR7 | - | - |
| | SRC | 2SRC | 1FMK | 4XI2 (39%) |
| | Thrombin | 1BA8 | 2AFQ | 1EAI (30%) |
