## Supplementary Information Table S2 for "PL-PatchSurfer3: Improved Structure-Based Virtual Screening for Structure Variation Using 3D Zernike Descriptors"

**Table S2**. Virtual screening results of individual targets in the DUD benchmark dataset. Dissimilarity indicates the average dissimilarity of active compounds from the cognate ligand calculated by SIMCOMP. PL-PatchSurfer2 results of all the targets were taken from the test set results.

**A**. PL-PatchSurfer3 (LCS)

| Target | Dissimilarity | $EF_{1\%}$ | $EF_{5\%}$ | $EF_{10\%}$ | AUC | BEDROC |
|---|---|---|---|---|---|---|
| AmpC | 0.754 | 0.00 | 0.00 | 0.95 | 0.513 | 0.056 |
| AR | 0.752 | 10.91 | 4.86 | 2.57 | 0.434 | 0.252 |
| CDK2 | 0.785 | 22.00 | 5.60 | 3.40 | 0.502 | 0.357 |
| COX-2 | 0.543 | 8.33 | 3.67 | 2.17 | 0.602 | 0.212 |
| EGFr | 0.576 | 20.00 | 4.00 | 2.17 | 0.506 | 0.264 |
| ER: antagonist | 0.579 | 8.18 | 4.14 | 5.13 | 0.813 | 0.310 |
| FXa | 0.761 | 5.00 | 2.00 | 1.83 | 0.432 | 0.147 |
| HIVRT | 0.769 | 20.00 | 5.00 | 3.50 | 0.613 | 0.337 |
| PARP | 0.716 | 23.33 | 5.51 | 3.03 | 0.531 | 0.339 |
| PPARγ | 0.314 | 1.25 | 1.49 | 1.36 | 0.493 | 0.091 |
| RXRα | 0.462 | 15.00 | 10.00 | 5.50 | 0.825 | 0.484 |
| VEGFr2 | 0.805 | 21.82 | 5.95 | 3.51 | 0.539 | 0.367 |
| AChE | 0.512 | 11.67 | 5.00 | 3.67 | 0.609 | 0.282 |
| COX-1 | 0.623 | 12.86 | 4.86 | 2.80 | 0.466 | 0.274 |
| ER: agonist | 0.760 | 13.50 | 5.70 | 3.43 | 0.616 | 0.319 |
| FGFr1 | 0.823 | 15.00 | 4.67 | 2.50 | 0.410 | 0.263 |
| GR | 0.885 | 10.43 | 3.33 | 2.05 | 0.627 | 0.211 |
| HIVPR | 0.804 | 20.00 | 9.87 | 6.23 | 0.802 | 0.551 |
| MR | 0.518 | 15.00 | 5.45 | 2.67 | 0.673 | 0.287 |
| NA | 0.669 | 10.71 | 4.52 | 2.45 | 0.566 | 0.258 |
| P38 MAP | 0.730 | 11.67 | 5.00 | 3.17 | 0.573 | 0.293 |
| PDGFrb | 0.699 | 16.67 | 4.67 | 2.67 | 0.382 | 0.278 |
| PR | 0.866 | 18.75 | 5.25 | 2.59 | 0.414 | 0.313 |
| SRC | 0.758 | 20.00 | 6.67 | 4.00 | 0.579 | 0.395 |
| Thrombin | 0.771 | 15.79 | 5.31 | 4.06 | 0.641 | 0.338 |

**B**. PL-PatchSurfer3 (BWS)

| Target | Dissimilarity | $EF_{1\%}$ | $EF_{5\%}$ | $EF_{10\%}$ | AUC | BEDROC |
|---|---|---|---|---|---|---|
| AmpC | 0.754 | 0.00 | 0.00 | 0.48 | 0.411 | 0.016 |
| AR | 0.752 | 9.55 | 5.41 | 3.11 | 0.619 | 0.280 |
| CDK2 | 0.785 | 24.00 | 6.00 | 3.60 | 0.507 | 0.381 |
| COX-2 | 0.543 | 11.67 | 5.00 | 3.83 | 0.700 | 0.311 |
| EGFr | 0.576 | 18.33 | 4.00 | 2.00 | 0.493 | 0.260 |
| ER: antagonist | 0.579 | 5.45 | 5.17 | 4.36 | 0.833 | 0.304 |
| FXa | 0.761 | 5.00 | 3.00 | 1.83 | 0.457 | 0.164 |
| HIVRT | 0.769 | 17.50 | 6.50 | 3.75 | 0.646 | 0.379 |
| PARP | 0.716 | 20.00 | 6.12 | 3.94 | 0.551 | 0.373 |
| PPARγ | 0.314 | 0.00 | 0.50 | 0.49 | 0.487 | 0.039 |
| RXRα | 0.462 | 15.00 | 9.00 | 6.00 | 0.894 | 0.499 |
| VEGFr2 | 0.805 | 21.82 | 6.49 | 3.65 | 0.546 | 0.393 |
| AChE | 0.512 | 10.00 | 4.00 | 2.83 | 0.588 | 0.257 |
| COX-1 | 0.623 | 21.43 | 4.86 | 2.80 | 0.464 | 0.304 |
| ER: agonist | 0.760 | 21.00 | 8.70 | 5.52 | 0.756 | 0.503 |
| FGFr1 | 0.823 | 15.00 | 4.67 | 2.67 | 0.479 | 0.269 |
| GR | 0.885 | 14.35 | 5.13 | 3.85 | 0.783 | 0.325 |
| HIVPR | 0.804 | 24.00 | 11.01 | 6.42 | 0.826 | 0.615 |
| MR | 0.518 | 15.00 | 5.45 | 2.67 | 0.826 | 0.306 |
| NA | 0.669 | 8.57 | 3.70 | 2.65 | 0.575 | 0.238 |
| P38 MAP | 0.730 | 11.67 | 3.67 | 2.50 | 0.601 | 0.234 |
| PDGFrb | 0.699 | 15.00 | 4.67 | 3.00 | 0.392 | 0.288 |
| PR | 0.866 | 18.75 | 6.00 | 3.70 | 0.472 | 0.364 |
| SRC | 0.758 | 20.00 | 6.00 | 4.33 | 0.601 | 0.396 |
| Thrombin | 0.771 | 15.79 | 5.21 | 3.34 | 0.606 | 0.313 |

**C**. AutoDock Vina

| Target | Dissimilarity | $EF_{1\%}$ | $EF_{5\%}$ | $EF_{10\%}$ | AUC | BEDROC |
|---|---|---|---|---|---|---|
| AmpC | 0.754 | 0.00 | 0.00 | 0.48 | 0.242 | 0.010 |
| AR | 0.752 | 16.36 | 10.00 | 6.49 | 0.771 | 0.539 |
| CDK2 | 0.785 | 8.00 | 6.40 | 4.00 | 0.665 | 0.322 |
| COX-2 | 0.543 | 25.00 | 12.67 | 6.83 | 0.834 | 0.673 |
| EGFr | 0.576 | 6.67 | 2.33 | 2.17 | 0.567 | 0.173 |
| ER: antagonist | 0.579 | 8.18 | 4.66 | 3.08 | 0.612 | 0.277 |
| FXa | 0.761 | 1.67 | 2.67 | 2.17 | 0.665 | 0.135 |
| HIVRT | 0.769 | 7.50 | 4.00 | 3.00 | 0.627 | 0.244 |
| PARP | 0.716 | 6.67 | 4.90 | 4.54 | 0.734 | 0.287 |
| PPARγ | 0.314 | 1.25 | 3.22 | 2.59 | 0.665 | 0.181 |
| RXRα | 0.462 | 25.00 | 15.00 | 8.00 | 0.924 | 0.747 |
| VEGFr2 | 0.805 | 8.18 | 3.51 | 2.70 | 0.544 | 0.216 |
| AChE | 0.512 | 5.00 | 3.33 | 3.33 | 0.724 | 0.230 |
| COX-1 | 0.623 | 12.86 | 8.11 | 5.60 | 0.711 | 0.438 |
| ER: agonist | 0.760 | 13.50 | 9.00 | 5.07 | 0.794 | 0.479 |
| FGFr1 | 0.823 | 0.00 | 0.00 | 0.00 | 0.457 | 0.017 |
| GR | 0.885 | 7.83 | 2.31 | 1.54 | 0.544 | 0.157 |
| HIVPR | 0.804 | 4.00 | 5.32 | 4.34 | 0.736 | 0.286 |
| MR | 0.518 | 22.50 | 13.64 | 7.33 | 0.800 | 0.694 |
| NA | 0.669 | 0.00 | 0.41 | 0.41 | 0.379 | 0.019 |
| P38 MAP | 0.730 | 0.00 | 2.00 | 1.67 | 0.595 | 0.117 |
| PDGFrb | 0.699 | 3.33 | 1.33 | 0.67 | 0.308 | 0.068 |
| PR | 0.866 | 0.00 | 1.50 | 1.11 | 0.427 | 0.060 |
| SRC | 0.758 | 3.33 | 4.33 | 3.00 | 0.682 | 0.217 |
| Thrombin | 0.771 | 11.05 | 5.62 | 4.22 | 0.729 | 0.320 |

**D**. DOCK6

| Target | Dissimilarity | $EF_{1\%}$ | $EF_{5\%}$ | $EF_{10\%}$ | AUC | BEDROC |
|---|---|---|---|---|---|---|
| AmpC | 0.754 | 20.00 | 4.84 | 2.38 | 0.595 | 0.299 |
| AR | 0.752 | 4.09 | 1.08 | 0.68 | 0.313 | 0.075 |
| CDK2 | 0.785 | 22.00 | 6.40 | 3.40 | 0.567 | 0.372 |
| COX-2 | 0.543 | 0.00 | 1.67 | 1.17 | 0.385 | 0.078 |
| EGFr | 0.576 | 18.33 | 4.67 | 2.67 | 0.512 | 0.305 |
| ER: antagonist | 0.579 | 16.36 | 4.66 | 3.33 | 0.616 | 0.308 |
| FXa | 0.761 | 8.33 | 2.67 | 1.83 | 0.605 | 0.194 |
| HIVRT | 0.769 | 5.00 | 3.00 | 1.75 | 0.442 | 0.158 |
| PARP | 0.716 | 16.67 | 5.51 | 3.33 | 0.548 | 0.332 |
| PPARγ | 0.314 | 0.00 | 1.49 | 0.86 | 0.418 | 0.066 |
| RXRα | 0.462 | 30.00 | 9.00 | 6.00 | 0.752 | 0.547 |
| VEGFr2 | 0.805 | 16.36 | 4.86 | 2.57 | 0.464 | 0.303 |
| AChE | 0.512 | 6.67 | 3.33 | 2.17 | 0.644 | 0.195 |
| COX-1 | 0.623 | 0.00 | 0.81 | 0.80 | 0.510 | 0.045 |
| ER: agonist | 0.760 | 10.50 | 3.90 | 2.09 | 0.492 | 0.208 |
| FGFr1 | 0.823 | 16.67 | 4.33 | 2.50 | 0.419 | 0.276 |
| GR | 0.885 | 5.22 | 1.28 | 0.90 | 0.368 | 0.097 |
| HIVPR | 0.804 | 2.00 | 1.90 | 1.70 | 0.280 | 0.121 |
| MR | 0.518 | 15.00 | 2.73 | 1.33 | 0.399 | 0.175 |
| NA | 0.669 | 23.57 | 12.33 | 6.73 | 0.827 | 0.629 |
| P38 MAP | 0.730 | 8.33 | 3.00 | 2.33 | 0.528 | 0.195 |
| PDGFrb | 0.699 | 5.00 | 1.67 | 0.83 | 0.244 | 0.099 |
| PR | 0.866 | 3.75 | 1.50 | 1.11 | 0.344 | 0.098 |
| SRC | 0.758 | 23.33 | 7.67 | 3.83 | 0.517 | 0.432 |
| Thrombin | 0.771 | 9.47 | 6.25 | 5.47 | 0.753 | 0.378 |
