## Supplementary Information Table S3 for "PL-PatchSurfer3: Improved Structure-Based Virtual Screening for Structure Variation Using 3D Zernike Descriptors"

**Table S3**. Virtual screening results for individual targets in the apo structure dataset. Cα-RMSD is a binding site residue Cα-RMSD between holo and apo forms.

**A**. PL-PatchSurfer3 (LCS)

| Targets | Cα-RMSD (Å) | $EF_{1\%}$ | $EF_{5\%}$ | $EF_{10\%}$ | AUC | BEDROC |
|---|---|---|---|---|---|---|
| AmpC | 0.17 | 0.00 | 0.00 | 0.47 | 0.338 | 0.023 |
| CDK2 | 0.30 | 20.00 | 6.40 | 3.80 | 0.502 | 0.367 |
| COX-2 | 0.38 | 6.67 | 1.33 | 1.67 | 0.452 | 0.123 |
| EGFr | 0.39 | 20.00 | 4.33 | 2.17 | 0.500 | 0.270 |
| ER: antagonist | 1.24 | 5.45 | 3.10 | 3.33 | 0.772 | 0.216 |
| FXa | 0.45 | 6.67 | 3.33 | 2.50 | 0.578 | 0.200 |
| PARP | 0.40 | 13.33 | 4.29 | 2.73 | 0.458 | 0.269 |
| PPARγ | 0.65 | 2.50 | 1.49 | 2.10 | 0.606 | 0.131 |
| RXRα | 3.77 | 15.00 | 3.00 | 3.00 | 0.569 | 0.240 |
| AChE | 0.27 | 10.00 | 3.67 | 2.83 | 0.599 | 0.234 |
| COX-1 | 0.30 | 8.57 | 4.05 | 2.80 | 0.456 | 0.238 |
| ER: agonist | 0.33 | 13.50 | 5.10 | 3.43 | 0.641 | 0.325 |
| FGFr1 | 0.19 | 13.33 | 4.33 | 2.33 | 0.430 | 0.256 |
| HIVPR | 2.08 | 12.00 | 7.97 | 4.72 | 0.703 | 0.412 |
| NA | 0.16 | 4.29 | 3.29 | 2.24 | 0.525 | 0.191 |
| P38 MAP | 1.85 | 5.00 | 3.33 | 1.83 | 0.423 | 0.165 |
| SRC | 2.14 | 16.67 | 6.67 | 4.17 | 0.561 | 0.380 |
| Thrombin | 1.38 | 20.53 | 6.88 | 4.84 | 0.679 | 0.433 |

**B**. PL-PatchSurfer3 (BWS)

| Targets | Cα-RMSD (Å) | $EF_{1\%}$ | $EF_{5\%}$ | $EF_{10\%}$ | AUC | BEDROC |
|---|---|---|---|---|---|---|
| AmpC | 0.17 | 0.00 | 0.00 | 0.48 | 0.299 | 0.013 |
| CDK2 | 0.30 | 26.00 | 7.20 | 3.60 | 0.504 | 0.415 |
| COX-2 | 0.38 | 6.67 | 3.33 | 2.33 | 0.568 | 0.177 |
| EGFr | 0.39 | 20.00 | 4.00 | 2.17 | 0.504 | 0.266 |
| ER: antagonist | 1.24 | 2.73 | 3.10 | 3.59 | 0.800 | 0.237 |
| FXa | 0.45 | 10.00 | 3.67 | 2.50 | 0.636 | 0.218 |
| PARP | 0.40 | 16.67 | 4.29 | 3.03 | 0.504 | 0.290 |
| PPARγ | 0.65 | 3.75 | 1.74 | 1.85 | 0.582 | 0.131 |
| RXRα | 3.77 | 15.00 | 6.00 | 4.00 | 0.682 | 0.347 |
| AChE | 0.27 | 10.00 | 4.33 | 2.83 | 0.594 | 0.237 |
| COX-1 | 0.30 | 21.43 | 4.05 | 2.40 | 0.449 | 0.274 |
| ER: agonist | 0.33 | 19.50 | 9.90 | 5.97 | 0.776 | 0.536 |
| FGFr1 | 0.19 | 15.00 | 4.67 | 2.67 | 0.503 | 0.270 |
| HIVPR | 2.08 | 16.00 | 8.73 | 4.91 | 0.709 | 0.471 |
| NA | 0.16 | 8.57 | 3.29 | 2.45 | 0.549 | 0.211 |
| P38 MAP | 1.85 | 6.67 | 2.67 | 1.67 | 0.451 | 0.170 |
| SRC | 2.14 | 18.33 | 6.33 | 4.00 | 0.589 | 0.386 |
| Thrombin | 1.38 | 20.53 | 7.19 | 5.16 | 0.694 | 0.451 |

**C**. AutoDock Vina

| Targets | Cα-RMSD (Å) | $EF_{1\%}$ | $EF_{5\%}$ | $EF_{10\%}$ | AUC | BEDROC |
|---|---|---|---|---|---|---|
| AmpC | 0.17 | 0.00 | 0.00 | 0.00 | 0.194 | 0.000 |
| CDK2 | 0.30 | 0.00 | 0.40 | 1.80 | 0.541 | 0.074 |
| COX-2 | 0.38 | 21.67 | 11.67 | 7.50 | 0.838 | 0.646 |
| EGFr | 0.39 | 5.00 | 4.00 | 2.67 | 0.626 | 0.216 |
| ER: antagonist | 1.24 | 0.00 | 0.00 | 0.00 | 0.428 | 0.005 |
| FXa | 0.45 | 3.33 | 3.00 | 1.83 | 0.643 | 0.164 |
| PARP | 0.40 | 6.67 | 2.45 | 2.42 | 0.606 | 0.182 |
| PPARγ | 0.65 | 1.25 | 0.25 | 0.86 | 0.584 | 0.067 |
| RXRα | 3.77 | 0.00 | 7.00 | 5.50 | 0.918 | 0.360 |
| AChE | 0.27 | 3.33 | 2.00 | 1.50 | 0.632 | 0.134 |
| COX-1 | 0.30 | 4.29 | 2.43 | 3.20 | 0.700 | 0.205 |
| ER: agonist | 0.33 | 12.00 | 7.80 | 4.18 | 0.742 | 0.400 |
| FGFr1 | 0.19 | 0.00 | 0.67 | 0.83 | 0.444 | 0.041 |
| HIVPR | 2.08 | 4.00 | 1.90 | 1.51 | 0.542 | 0.113 |
| NA | 0.16 | 2.14 | 0.41 | 0.20 | 0.363 | 0.029 |
| P38 MAP | 1.85 | 0.00 | 0.00 | 0.00 | 0.222 | 0.001 |
| SRC | 2.14 | 0.00 | 1.33 | 1.00 | 0.351 | 0.069 |
| Thrombin | 1.38 | 4.74 | 2.19 | 1.72 | 0.568 | 0.151 |

**D**. DOCK6

| Targets | Cα-RMSD (Å) | $EF_{1\%}$ | $EF_{5\%}$ | $EF_{10\%}$ | AUC | BEDROC |
|---|---|---|---|---|---|---|
| AmpC | 0.17 | 20.00 | 4.84 | 2.38 | 0.581 | 0.284 |
| CDK2 | 0.30 | 12.00 | 4.00 | 2.20 | 0.468 | 0.224 |
| COX-2 | 0.38 | 3.33 | 2.33 | 2.00 | 0.498 | 0.138 |
| EGFr | 0.39 | 16.67 | 3.67 | 2.50 | 0.441 | 0.253 |
| ER: antagonist | 1.24 | 0.00 | 0.52 | 0.51 | 0.512 | 0.039 |
| FXa | 0.45 | 1.76 | 2.36 | 1.84 | 0.526 | 0.142 |
| PARP | 0.40 | 16.67 | 6.12 | 3.33 | 0.586 | 0.336 |
| PPARγ | 0.65 | 7.50 | 5.21 | 5.06 | 0.792 | 0.343 |
| RXRα | 3.77 | 0.00 | 0.00 | 0.50 | 0.538 | 0.042 |
| AChE | 0.27 | 3.33 | 1.67 | 1.33 | 0.437 | 0.111 |
| COX-1 | 0.30 | 4.29 | 2.43 | 2.00 | 0.418 | 0.160 |
| ER: agonist | 0.33 | 7.50 | 4.50 | 2.99 | 0.549 | 0.245 |
| FGFr1 | 0.19 | 15.00 | 4.67 | 2.67 | 0.391 | 0.279 |
| HIVPR | 2.08 | 0.00 | 1.14 | 1.13 | 0.247 | 0.071 |
| NA | 0.16 | 12.86 | 9.86 | 6.33 | 0.840 | 0.526 |
| P38 MAP | 1.85 | 6.67 | 2.00 | 1.33 | 0.388 | 0.133 |
| SRC | 2.14 | 5.00 | 3.67 | 2.33 | 0.421 | 0.196 |
| Thrombin | 1.38 | 11.05 | 8.13 | 6.25 | 0.829 | 0.458 |
