## Supplementary Information Table S4 for "PL-PatchSurfer3: Improved Structure-Based Virtual Screening for Structure Variation Using 3D Zernike Descriptors"

**Table S4**. Virtual screening results for individual targets in the template-based model dataset. RMSD is calculated after the target structure and the model superimposed by TM-align.

**A**. PL-PatchSurfer3 (LCS)

| Targets | Seq. Id. with Templates (%) | $EF_{1\%}$ | $EF_{5\%}$ | $EF_{10\%}$ | AUC | BEDROC |
|---|---|---|---|---|---|---|
| AmpC | 40 | 0.00 | 0.97 | 0.95 | 0.514 | 0.073 |
| CDK2 | 30 | 12.00 | 5.20 | 3.00 | 0.475 | 0.292 |
| COX-2 | 64 | 5.00 | 1.00 | 0.67 | 0.341 | 0.083 |
| EGFr | 78 | 15.00 | 4.33 | 2.50 | 0.526 | 0.268 |
| ER: antagonist | 30 | 5.45 | 4.14 | 3.08 | 0.728 | 0.225 |
| FXa | 30 | 5.00 | 2.33 | 2.00 | 0.576 | 0.161 |
| PARP | 45 | 10.00 | 6.37 | 2.12 | 0.409 | 0.202 |
| PPARγ | 30 | 1.25 | 1.24 | 1.73 | 0.626 | 0.096 |
| RXRα | 30 | 10.00 | 5.00 | 4.00 | 0.721 | 0.312 |
| AChE | 30 | 8.33 | 6.00 | 3.83 | 0.609 | 0.327 |
| COX-1 | 64 | 12.86 | 5.68 | 2.80 | 0.479 | 0.304 |
| ER: agonist | 30 | 9.00 | 5.10 | 3.43 | 0.637 | 0.299 |
| FGFr1 | 32 | 16.67 | 4.00 | 2.33 | 0.393 | 0.246 |
| HIVPR | 30 | 16.00 | 9.11 | 5.28 | 0.748 | 0.464 |
| NA | 31 | 10.71 | 2.05 | 1.84 | 0.475 | 0.171 |
| P38 MAP | 35 | 8.33 | 3.33 | 2.17 | 0.443 | 0.204 |
| SRC | 39 | 15.00 | 5.33 | 3.00 | 0.540 | 0.311 |
| Thrombin | 30 | 14.21 | 5.63 | 4.22 | 0.657 | 0.363 |

**B**. PL-PatchSurfer3 (BWS)

| Targets | Seq. Id. with Templates (%) | $EF_{1\%}$ | $EF_{5\%}$ | $EF_{10\%}$ | AUC | BEDROC |
|---|---|---|---|---|---|---|
| AmpC | 40 | 0.00 | 0.97 | 0.48 | 0.480 | 0.039 |
| CDK2 | 30 | 20.00 | 6.00 | 3.40 | 0.500 | 0.374 |
| COX-2 | 64 | 5.00 | 2.33 | 1.17 | 0.441 | 0.129 |
| EGFr | 78 | 16.67 | 4.00 | 2.50 | 0.530 | 0.265 |
| ER: antagonist | 30 | 2.73 | 4.14 | 4.10 | 0.792 | 0.269 |
| FXa | 30 | 6.67 | 3.00 | 2.17 | 0.559 | 0.181 |
| PARP | 45 | 10.00 | 4.29 | 2.72 | 0.430 | 0.261 |
| PPARγ | 30 | 0.00 | 0.74 | 1.23 | 0.597 | 0.071 |
| RXRα | 30 | 10.00 | 7.00 | 5.00 | 0.787 | 0.404 |
| AChE | 30 | 10.00 | 5.00 | 3.83 | 0.616 | 0.331 |
| COX-1 | 64 | 21.43 | 4.05 | 2.40 | 0.467 | 0.281 |
| ER: agonist | 30 | 15.00 | 8.40 | 5.22 | 0.763 | 0.482 |
| FGFr1 | 32 | 13.33 | 4.00 | 2.33 | 0.480 | 0.251 |
| HIVPR | 30 | 20.00 | 9.49 | 5.66 | 0.750 | 0.521 |
| NA | 31 | 8.57 | 3.29 | 1.84 | 0.495 | 0.183 |
| P38 MAP | 35 | 10.00 | 4.00 | 2.17 | 0.437 | 0.208 |
| SRC | 39 | 11.67 | 5.67 | 3.33 | 0.587 | 0.309 |
| Thrombin | 30 | 14.21 | 6.25 | 3.91 | 0.660 | 0.369 |

**C**. AutoDock Vina

| Targets | Seq. Id. with Templates (%) | $EF_{1\%}$ | $EF_{5\%}$ | $EF_{10\%}$ | AUC | BEDROC |
|---|---|---|---|---|---|---|
| AmpC | 40 | 0.00 | 0.00 | 0.00 | 0.208 | 0.000 |
| CDK2 | 30 | 0.00 | 0.40 | 0.40 | 0.462 | 0.022 |
| COX-2 | 64 | 1.67 | 0.67 | 0.83 | 0.539 | 0.048 |
| EGFr | 78 | 1.67 | 3.67 | 2.67 | 0.666 | 0.197 |
| ER: antagonist | 30 | 0.00 | 0.00 | 0.00 | 0.393 | 0.006 |
| FXa | 30 | 0.00 | 3.33 | 3.50 | 0.705 | 0.201 |
| PARP | 45 | 13.33 | 7.96 | 5.15 | 0.573 | 0.442 |
| PPARγ | 30 | 0.00 | 0.74 | 1.11 | 0.660 | 0.064 |
| RXRα | 30 | 0.00 | 0.00 | 0.50 | 0.578 | 0.045 |
| AChE | 30 | 6.67 | 1.67 | 1.17 | 0.533 | 0.124 |
| COX-1 | 64 | 0.00 | 0.00 | 0.00 | 0.453 | 0.011 |
| ER: agonist | 30 | 4.50 | 2.70 | 2.84 | 0.654 | 0.185 |
| FGFr1 | 32 | 0.00 | 0.67 | 0.83 | 0.434 | 0.049 |
| HIVPR | 30 | 4.00 | 1.90 | 1.13 | 0.597 | 0.113 |
| NA | 31 | 0.00 | 0.00 | 0.00 | 0.169 | 0.001 |
| P38 MAP | 35 | 1.67 | 0.33 | 0.17 | 0.281 | 0.022 |
| SRC | 39 | 0.00 | 0.00 | 0.33 | 0.284 | 0.011 |
| Thrombin | 30 | 0.00 | 0.94 | 1.09 | 0.417 | 0.066 |

**D**. DOCK6

| Targets | Seq. Id. with Templates (%) | $EF_{1\%}$ | $EF_{5\%}$ | $EF_{10\%}$ | AUC | BEDROC |
|---|---|---|---|---|---|---|
| AmpC | 40 | 10.00 | 5.81 | 3.33 | 0.592 | 0.294 |
| CDK2 | 30 | 6.00 | 2.40 | 2.60 | 0.501 | 0.183 |
| COX-2 | 64 | 3.33 | 1.67 | 0.83 | 0.356 | 0.087 |
| EGFr | 78 | 16.67 | 3.67 | 2.00 | 0.456 | 0.237 |
| ER: antagonist | 30 | 0.00 | 0.52 | 0.26 | 0.302 | 0.016 |
| FXa | 30 | 5.00 | 4.33 | 3.50 | 0.686 | 0.246 |
| PARP | 45 | 16.67 | 5.51 | 3.03 | 0.476 | 0.330 |
| PPARγ | 30 | 1.25 | 2.23 | 1.60 | 0.627 | 0.124 |
| RXRα | 30 | 0.00 | 0.00 | 0.00 | 0.573 | 0.014 |
| AChE | 30 | 1.67 | 1.00 | 1.17 | 0.492 | 0.081 |
| COX-1 | 64 | 0.00 | 0.00 | 0.00 | 0.300 | 0.000 |
| ER: agonist | 30 | 3.00 | 1.20 | 1.04 | 0.377 | 0.080 |
| FGFr1 | 32 | 6.67 | 2.33 | 1.83 | 0.377 | 0.149 |
| HIVPR | 30 | 0.00 | 1.14 | 0.75 | 0.238 | 0.050 |
| NA | 31 | 12.86 | 7.40 | 4.69 | 0.715 | 0.404 |
| P38 MAP | 35 | 18.33 | 6.00 | 3.67 | 0.529 | 0.350 |
| SRC | 39 | 16.67 | 6.00 | 3.50 | 0.460 | 0.352 |
| Thrombin | 30 | 6.32 | 4.06 | 3.28 | 0.651 | 0.253 |
