## Supplementary Information Table S5 for "PL-PatchSurfer3: Improved Structure-Based Virtual Screening for Structure Variation Using 3D Zernike Descriptors"

**Table S5**. Virtual screening results of individual targets in the AlphaFold2 benchmark dataset.

**A**. PL-PatchSurfer3 (LCS)

| Target | $EF_{1\%}$ | $EF_{5\%}$ | $EF_{10\%}$ | AUC | BEDROC |
|---|---|---|---|---|---|
| CDK2 | 14.00 | 4.80 | 2.80 | 0.498 | 0.288 |
| EGFr | 20.00 | 4.00 | 2.17 | 0.483 | 0.262 |
| FGFr1 | 15.00 | 4.33 | 2.17 | 0.410 | 0.262 |
| P38 MAP | 6.67 | 3.67 | 2.33 | 0.499 | 0.204 |
| SRC | 16.67 | 7.00 | 4.00 | 0.548 | 0.373 |
| VEGFr2 | 20.45 | 5.68 | 3.78 | 0.534 | 0.359 |

**B**. PL-PatchSurfer3 (BWS)

| Target | $EF_{1\%}$ | $EF_{5\%}$ | $EF_{10\%}$ | AUC | BEDROC |
|---|---|---|---|---|---|
| CDK2 | 20.00 | 6.00 | 4.00 | 0.517 | 0.379 |
| EGFr | 16.67 | 4.33 | 2.50 | 0.476 | 0.274 |
| FGFr1 | 15.00 | 4.33 | 2.33 | 0.472 | 0.269 |
| P38 MAP | 8.33 | 3.00 | 2.00 | 0.506 | 0.190 |
| SRC | 18.33 | 6.00 | 4.00 | 0.588 | 0.356 |
| VEGFr2 | 19.09 | 5.95 | 3.92 | 0.541 | 0.367 |

**C**. AutoDock Vina

| Target | $EF_{1\%}$ | $EF_{5\%}$ | $EF_{10\%}$ | AUC | BEDROC |
|---|---|---|---|---|---|
| CDK2 | 2.00 | 4.00 | 3.60 | 0.621 | 0.221 |
| EGFr | 1.67 | 2.00 | 1.33 | 0.576 | 0.103 |
| FGFr1 | 0.00 | 0.67 | 1.33 | 0.482 | 0.063 |
| P38 MAP | 0.00 | 0.33 | 1.67 | 0.584 | 0.063 |
| SRC | 0.00 | 2.00 | 2.50 | 0.624 | 0.142 |
| VEGFr2 | 13.64 | 4.32 | 2.84 | 0.584 | 0.281 |

**D**. DOCK6

| Target | $EF_{1\%}$ | $EF_{5\%}$ | $EF_{10\%}$ | AUC | BEDROC |
|---|---|---|---|---|---|
| CDK2 | 16.00 | 5.60 | 3.20 | 0.565 | 0.332 |
| EGFr | 13.33 | 4.00 | 2.17 | 0.414 | 0.244 |
| FGFr1 | 6.67 | 3.00 | 2.33 | 0.378 | 0.182 |
| P38 MAP | 3.33 | 2.67 | 1.83 | 0.514 | 0.136 |
| SRC | 10.00 | 4.33 | 3.67 | 0.490 | 0.293 |
| VEGFr2 | 20.45 | 5.14 | 2.84 | 0.472 | 0.341 |
